# Altered susceptibility of rare yeast species causing vulvovaginal candidiasis at acidic vaginal environment

**DOI:** 10.1101/2025.06.10.659005

**Authors:** Fang Liu, Yanxia Sun, Rongfeng Zhao, lu Zhang, Shuwen Deng

## Abstract

This study evaluated potency of antifungals against the rare emerging yeast causing VVC under the influence of acid pH (4.5). The antifungal susceptibility was evaluated by CLSI-M27A4 for 25 isolates represented 11 rare yeast species recovered from VVC patients. Among 11 yeast species (25 isolates), *Clavispora lusitaniae* (n=9) was the most common species, followed by *Cyberlindnera fabianii* (n=4)*, Kodamaea ohmeri, Candida metapsilosis* and *Candida bracarensis* (n=2, respectively)*, Pichia norvegensis, Kluyveromyces marxianus, Diutina rugosa, Tricholoma matsutake, Candida nivariensis* and *Candida africana* (n=1, respectively). The antifungal agents most affected by a lower pH were terconazole (4-fold higher, MIC >0.5-8mg/L) to all 11 yeast species; followed by amphotericin B (3-fold higher, MIC >2mg/L) to 9 species except *K. ohmeri* and *D. rugose*; nystatin (2-fold higher, MIC >2 - 4 mg/L) to 9 species except *C. nivariensis and C*.*africana*; micafungin (2-fold higher, MIC >0.125 - 1 mg/L) to 8 species except *C. metapsilosis, D. rugosa* and *Tricholoma matsutake* by acid pH, respectively. 5-flucytosine was not affected by pH to all tested yeast species, and maintained activity at low PH.

The rare yeast species tested shared high MICs of terconazole, amphotericin B, nystatin at pH4.5 compared with those at pH7.0. The *in vitro* antifungal susceptibility at pH 7.0 did not consistently predict the activity of antifungals against those rare yeast species at vaginal acidic pH. Further investigations assessing the clinical utility of antifungal susceptibility testing should take these findings into consideration.

## Introduction

Vulvovaginal candidiasis (VVC) affects millions of women annually. Although it represents a problem of global importance in public health, its exact incidence is unknown. Approximately one-third of vaginitis cases are caused by *Candida* (1).

VVC is most commonly caused by *Candida albicans* but can also be caused by non-*albicans Candida* species, including *Nakaseomyces glabrata, C. parapsilosis, C. tropicalis, P. kudriavzevii* and others. However, rare yeast other than the common *Candida* spp. have emerged as significant pathogens of VVC (2, 3). Whether this is due to improved diagnostics or whether there is a factual rise of uncommon yeasts emerging in the clinic, or both, remains unclear. Based on recent nomenclature and taxonomy, some species were moved to different genera, such as *Diutina rugose* (synonym: *Candida rugosa*), *Candida pararugosa, Trichomonascus ciferrii* (synonym: *Candida ciferrii*), *Candida inconspicua* (basionym: *Torulopsis inconspicua*) and *Pichia norvegensis* (synonym: *Candida norvegensis*).

These fungi are commonly encountered in the environment and frequent colonizers of human skin and mucosal surfaces (4, 5). These rare yeast occur in low percentage of VVC (<0.01 %), but is highly polyphyletic. Infections by these agents are hard to manage owing to limited clinical experience, lack of susceptibility data (6).

Unfortunately, the epidemiology of many of these rare and emerging infections is still not well studied, particularly with VVC (6, 7). Epidemiological cut-off values (ECOFFs) and clinical breakpoints (CBPs) for the international standard methods of CLSI and EUCAST have not yet been established for the rare yeast species to majority of antifungal drugs (16,17). A few case reports pointed to elevated MICs of at least one azole or of whole drug classes, such as in *T. ciferrii* (6), *C. guilliermondii, C. haemulonii, C. inconspicua* and *P. norvegensis* (4, 7). Echinocandins resistance was described in *T. ciferrii* (6).

Moreover, due to the normal pH of the vagina is 4 to 4.5, which remains unchange during VVC (8). Previous studies have found that some *Candida species* that are susceptible to antifungals *in vitro* (at pH 7.0) may be resistant to the same drug at acidic pH of vaginal environment (9). Mark Spitzer(10) measured at pH 7.0 and Ph 4.5 for fluconazole, itraconazole, miconazole, clotrimazole, terconazole, and nystatin to10 yeast species isolated from VVC, found that antifungals reduced *in vitro* potency when tested at lower pH, particularly for treatment of recurrent *N. glabrata* vaginitis, clinicians should recognize the limitations of *in vitro* susceptibility testing at pH 7.0. However, the most studies focused on the antifungal potency against the common yeast species (e.g. *C. albicans or N. glabrata*) caused VVC at acidic vaginal environment. Few data are available on rare yeast.

This study is to investigate antifungal susceptibility of the rare emerging yeast causing VVC to 13 antifungals against used in gynecological routines and echinocandins at acidic vaginal environment, to determine the extent to which this reduced effectiveness may contribute to clinical failure of therapy

## Materials and Methods

### Isolates and identification

A total of 25 rare yeast isolates were recovered from patients with vulvovaginal candidiasis at the gynecology clinic of the People’s hospital of Suzhou New District during Jan to Dec. 2021. The information on case definition, Patient data and clinical information, vaginal samples collecting have been reported in previous publication (3). The study was conducted in accordance with the Declaration of Helsinki. Ethical approval for the study was obtained from the Ethics Committee of the People’s Hospital of Suzhou New District (The IRB ethics number: 2022023). All participants signed an informed consent to the investigation.

All isolates were identified to the species level by sequencing the D1/D2 domain of 26S rDNA as described previously (3). GenBank accession numbers of D1/D2 sequences for 26 isolates studied are listed in table S1.

### Antifungal Susceptibility Testing

All isolates were tested for *in vitro* susceptibility to 13 antifungal drugs according to the CLSI M27A4/M44-ed3(11, 12). Antifungal drugs including fluconazole (FLC), itraconazole (ITR), voriconazole (VRC) and posaconazole (POS), terconazole (TEC), miconazole (MIC), clotrimazole (CLO), amphotericin B (AmB), 5-flucytosine (5-FC), nystatin (NYS), anidulafungin (ANF), caspofungin (CAS), micafungin (MFG) were obtained from Sigma Aldrich. *C. parapsilosis* ATCC 22019 and *P. kudriavzevii* ATCC 6258 were used as control strains. Colonies were suspended in sterile saline, and the final inoculum concentration of the suspension was adjusted to 0.5–2.5×10^3^ CFU/ mL.

The minimum inhibitory concentrations (MIC) at both pH 7.0 and pH 4.5 were read after 24 hours. MIC range, GMs, MIC_50/90_ were calculated for each species when testing ≥4 isolates, or range of MICs if < 4 isolates.

Clinical breakpoints (CBPs) or ECV of *C. albicans* set by CLSI served as a comparator for: *D. rugosa, K. marxianus, C. africana*; CBPs of *N. glabrata* served as a comparator for *C. nivariensis* and *C. bracarensis;* CBPs of *P. kudriavzevii* (= *Candida krusei*) for *P. norvegensis* as these species are phylogenetically closely related to the respective species (13). The interpretive breakpoints used in this study were listed in Table S2

To easy terminology, isolates with higher MIC values than the CBP or ECV of the comparative species are referred as resistant, and species with lower MIC values than the CBP or ECV as susceptible. Essential agreement (EA) was defined as a maximum difference of two 2-fold dilution steps between two MIC values (13).

## Results

### Identification of rare yeast at species level

In our study, 25 isolates of rare yeast recovered from the vaginal samples were identified into 11 species based on D1/D2 gene sequencing. *Cl. lusitaniae* was the most common species, followed by *Cy. fabianii* (n=4), *Kodamaea ohmeri, C. metapsilosis* and *C. bracarensis* (n=2, respectively), *P. norvegensis, Kl. marxianus(C. kefyr), Diutina rugosa, Tricholoma matsutake, C. nivariensis* and *C. africana* (n=1, respectively) (Fig.1).

**FIG 1:**
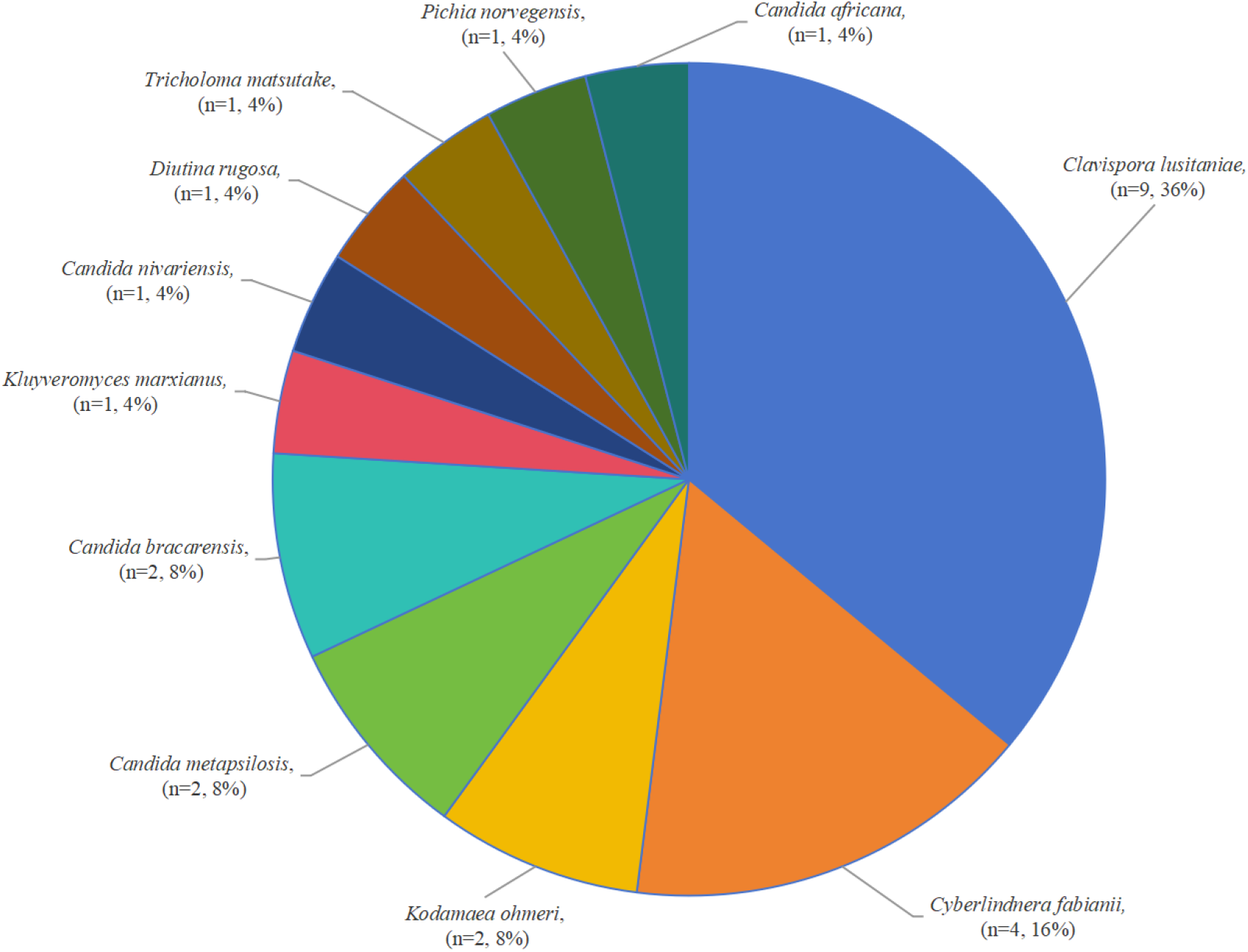
The distribution of 25 rare yeast species recovered from VVC.

### Antifungal susceptibility

Table 1 summarized the MICs and MIC range of 7 azoles to 25 rare yeast isolates at pH7.0 and pH4.5, respectively. Fig. 2 showed MIC Geometric means (GMs) of 7 azoles to rare yeast species (n≥2 isolates) at pH7.0 and pH4.5, respectively. Table S3 summarized MIC_50/90_, Geometric means (GMs) of 13 antifungals tested to rare yeast species (n≥2 isolates) at pH7.0 and pH4.5

**TABLE 1:** MICs and range of MIC of 7 azoles to 25 rare yeast isolates at pH7.0 and pH4.5.

| species (n) | pH | MIC rang (mg/L) |  |  |  |  |  |  |
| --- | --- | --- | --- | --- | --- | --- | --- | --- |
|  |  | fluconazole | voriconazole | itraconazole | posaconazole | terconazole | miconazole | clotrimazole |
| *Clavispora lusitanae*<br>n=9 | 4.5 | 0.125-2 | 0.031 | 0.063-0.5 | 0.063-0.125 | 1-2 | 0.031-0.063 | 0.031-0.063 |
|  | 7.0 | 0.125-1 | 0.031 | 0.031-0.063 | 0.031 | 0.031-0.063 | 0.031-0.063 | 0.031-0.063 |
| *Cyberlindnera fabianii*<br>n=4 | 4.5 | 0.5-4 | 0.031-0.125 | 0.5-1 | 0.5-1 | 4-8 | 0.25-0.5 | 0.25-0.5 |
|  | 7.0 | 1-8 | 0.031-0.125 | 0.25-0.5 | 0.25-1 | 0.125-0.5 | 1-2 | 0.25-1 |
| *Kodamaea ohmeri*<br>n=2 | 4.5 | 2-16 | 0.031-0.25 | 0.063-0.25 | 0.125-0.25 | 2-8 | 0.125-0.25 | 0.125-0.25 |
|  | 7 | 2-8 | 0.031-0.5 | 0.063-0.25 | 0.125-0.25 | 0.063-0.25 | 0.063-0.25 | 0.063-0.25 |
| *Candida metapsilosis*<br>n=2 | 4.5 | 1-1 | 0.031-0.031 | 0.125-0.125 | 0.125 | 1-4 | 0.063-0.125 | 0.031-0.031 |
|  | 7.0 | 0.5-1 | 0.031-0.031 | 0.031-0.063 | 0.031 | 0.031-0.063 | 0.25-0.25 | 0.031-0.031 |
| *Candida bracarensis*<br>n=2 | 4.5 | 4-8 | 0.125-0.25 | 1-1 | 1-1 | 4-8 | 0.125-0.125 | 0.5-0.5 |
|  | 7.0 | 1-2 | 0.031-0.063 | 0.063-0.125 | 0.063-0.125 | 0.031-0.063 | 0.031-0.063 | 0.031-0.125 |
| *Candida nivariensis*<br>n=1 | 4.5 | 4 | 0.125 | 0.5 | 0.5 | 4 | 0.125 | 0.25 |
|  | 7.0 | 1 | 0.031 | 0.063 | 0.063 | 0.031 | 0.031 | 0.031 |
| *Candida africana*<br>n=1 | 4.5 | 0.125 | 0.031 | 0.125 | 0.063 | 0.5 | 0.031 | 0.031 |
|  | 7.0 | 0.25 | 0.031 | 0.031 | 0.031 | 0.031 | 0.031 | 0.031 |
| *Kluyveromyces marxianus*<br>n=1 | 4.5 | 0.125 | 0.031 | 0.031 | 0.031 | 0.25 | 0.031 | 0.031 |
|  | 7.0 | 0.25 | 0.031 | 0.063 | 0.063 | 0.031 | 0.031 | 0.031 |
| *Diutina rugosa*<br>n=1 | 4.5 | 1 | 0.031 | 0.031 | 0.063 | 2 | 0.031 | 0.063 |
|  | 7.0 | 2 | 0.031 | 0.031 | 0.031 | 0.063 | 0.063 | 0.125 |
| *Tricholoma matsutake*<br>n=1 | 4.5 | 1 | 0.031 | 0.25 | 0.25 | 2 | 0.125 | 0.063 |
|  | 7.0 | 1 | 0.031 | 0.031 | 0.031 | 0.063 | 0.25 | 0.031 |
| *Pichia norvegensis*<br>n=1 | 4.5 | 32 | 0.5 | 0.5 | 0.25 | 8 | 1 | 0.25 |
|  | 7.0 | 8 | 0.5 | 0.5 | 0.25 | 0.5 | 4 | 0.25 |

**FIG 2:**
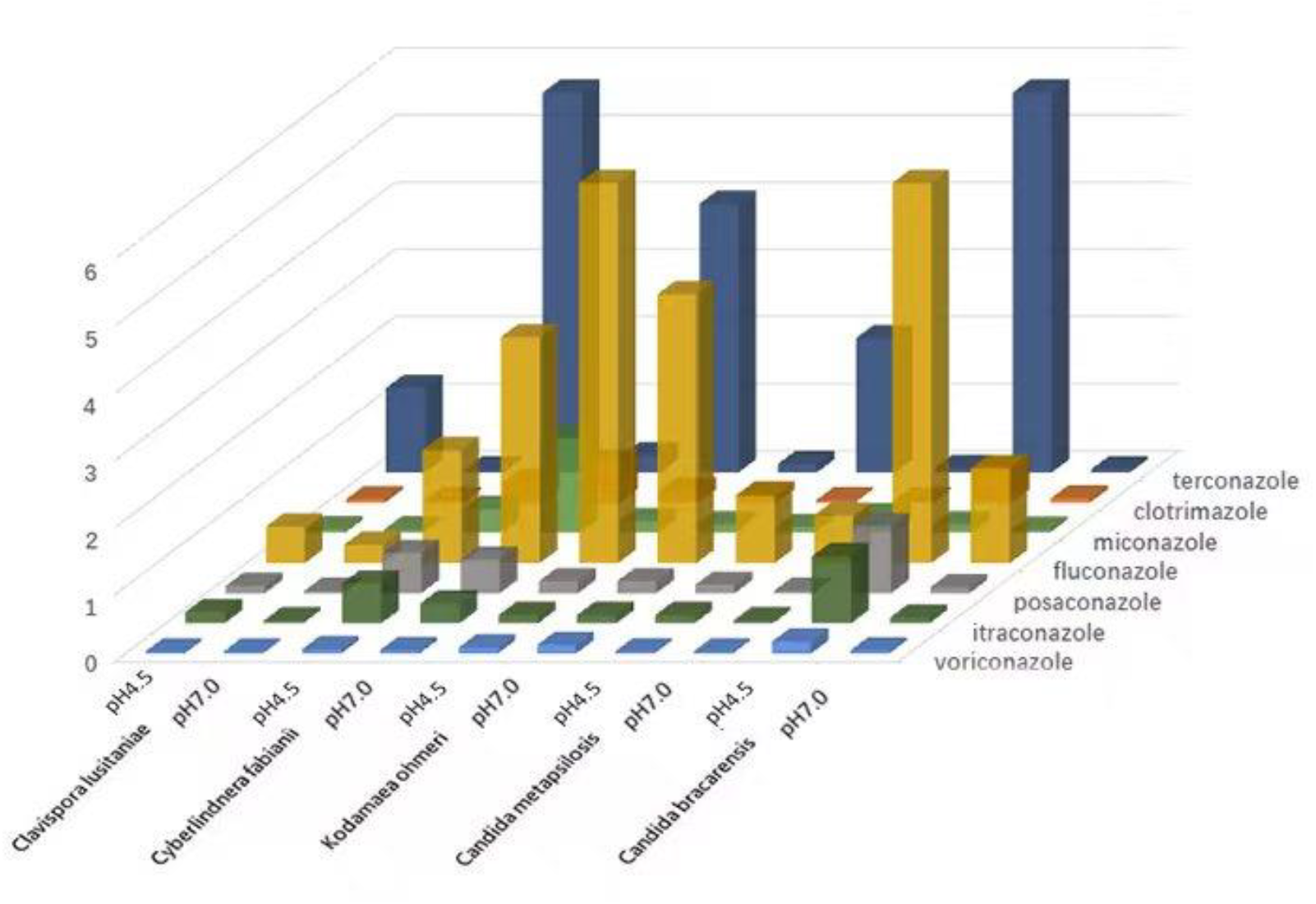
Geometric means (GMs) of 7 azoles to rare yeast species (n≥2 isolates) at pH7.0 and pH4.5

The most dramatic changes in MIC values between pH7.0 and pH4.5 were observed for terconazole, which the most species had > four 2-fold dilution steps increasing of MIC (>2mg/L) at pH4.5 when compared with those at pH7.0.

*C. bracarensis*(n=2) and *C. nivariensis*(n=1)in complex of *N. glabrata* were the species affected by the lower pH to all 7 azoles tested, with an increase in MIC > two 2-fold dilution steps at pH4.5 compared with those at pH7.0. However, these isolates remained susceptible to azoles with the increased MIC at pH4.5 when compared by CBP or ECV of *N. glabrata* (Table S2).

For *P. norvegensis*(n=1), a dramatic increases in MIC were seen for fluconazole only (32mg/L in pH4.0 and 8mg/L at pH7.0) besides of terconazole (8 mg/L in pH4.5/ 0.5 mg/L at pH7.0) at pH4.5. The other 5 azoles tested to this one isolate of *P. norvegensis* had unchanged MIC, and showing less susceptibility to all azoles at both pH levels.

The remaining species were unaffected by the alterations in pH except terconazole, which showed sensitive to all azoles tested at both pH levels.

Table 2 summarized MICs and MIC range of 3 echinocandins, amphotericin B, nystatin and 5-flucytosine to 25 rare yeast isolates at pH7.0 and pH4.5, respectively. Fig.3 showed MIC Geometric means (GMs) of 3 echinocandins, amphotericin B, nystatin and 5-flucytosine to rare yeast species (n≥2 isolates) at pH7.0 and pH4.5, respectively (Table S3).

**TABLE 2:** MICs and range of 3 echinocandins, amphotericin B, nystatin and 5-flucytosine to 25 rare yeast isolates at pH7.0 and pH4.5.

| species (n) | pH | MIC rang (mg/L) |  |  |  |  |  |
| --- | --- | --- | --- | --- | --- | --- | --- |
|  |  | anidulafungin | micafungin | caspofungin | amphotericin B | nystatin | 5-flucytosine |
| *Clavispora lusitaniae*<br>n=9 | 4.5 | 0.063-0.5 | 0.25-1 | 0.5-1 | 4 | 4-8 | 0.031 |
|  | 7.0 | 0.063-0.25 | 0.031-0.25 | 0.5-1 | 0.125-0.5 | 1-2 | 0.031 |
| *Cyberlindnera fabianii*<br>n=4 | 4.5 | 0.063-0.125 | 0.25-1 | 0.125-0.5 | 2-4 | 4-8 | 0.031 |
|  | 7.0 | 0.063-0.25 | 0.031-0.063 | 0.125-0.5 | 0.5-1 | 1-2 | 0.031-0.063 |
| *Kodamaea ohmeri*<br>n=2 | 4.5 | 0.125-0.125 | 0.25-0.5 | 0.25-0.25 | 0.25-0.5 | 2-8 | 0.031-0.031 |
|  | 7.0 | 0.25-0.25 | 0.063-0.125 | 0.25-0.5 | 0.5 | 1 | 0.031-0.063 |
| *Candida metapsilosis*<br>n=2 | 4.5 | 0.125-0.25 | 0.25-1 | 0.25-1 | 4-4 | 1-1 | 0.031-0.031 |
|  | 7.0 | 0.5-1 | 0.25-1 | 0.5-1 | 1-1 | 4-4 | 0.031-0.063 |
| *Candida bracarensis*<br>n=2 | 4.5 | 0.063-0.25 | 0.125-0.5 | 0.5-0.5 | 2-4 | 4-8 | 0.031-0.031 |
|  | 7.0 | 0.063-0.125 | 0.016-0.016 | 0.5-0.5 | 0.5-0.5 | 1-2 | 0.063-0.125 |
| *Candida nivariensis*<br>n=1 | 4.5 | 0.125 | 0.5 | 0.5 | 4 | 4 | 0.031 |
|  | 7.0 | 0.063 | 0.016 | 0.25 | 0.5 | 2 | 0.063 |
| *Candida africana*<br>n=1 | 4.5 | 0.031 | 0.125 | 0.5 | 4 | 2 | 0.031 |
|  | 7.0 | 0.016 | 0.031 | 0.25 | 0.25 | 2 | 0.031 |
| *Kluyveromyces marxianus*<br>n=1 | 4.5 | 0.5 | 0.25 | 0.063 | 2 | 4 | 0.031 |
|  | 7.0 | 0.125 | 0.063 | 0.125 | 0.5 | 1 | 0.031 |
| *Diutina rugosa*<br>n=1 | 4.5 | 0.5 | 0.125 | 2 | 2 | 8 | 0.031 |
|  | 7.0 | 1 | 0.25 | 2 | 2 | 4 | 0.063 |
| *Tricholoma matsutake*<br>n=1 | 4.5 | 0.25 | 1 | 0.5 | 4 | 4 | 0.031 |
|  | 7.0 | 0.5 | 0.5 | 0.5 | 0.5 | 1 | 0.031 |
| *Pichia norvegensis*<br>n=1 | 4.5 | 0.125 | 1 | 0.5 | 8 | 8 | 0.031 |
|  | 7.0 | 0.016 | 0.031 | 0.125 | 1 | 0.5 | 0.125 |

**FIG 3:**
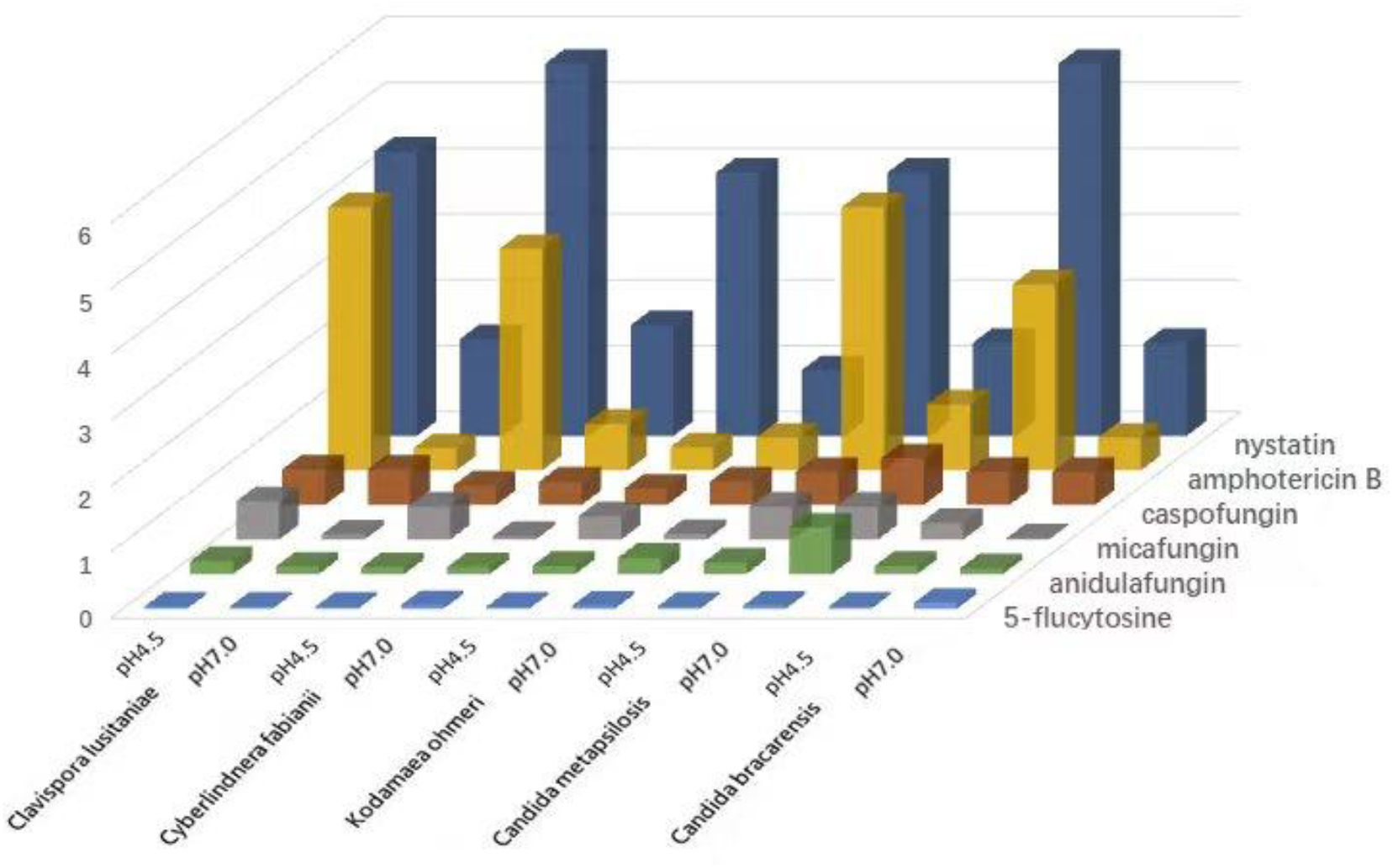
Geometric means (GMs) of 3 echinocandins, amphotericin B, nystatin and 5-flucytosine to rare yeast species (n≥2 isolates) at pH7.0 and pH4.5

Among 3 echinocandins tested in this study, micafungin was the most frequently affected by lower pH against 7 species (*Cl. lusitaniae, C. fabianii, K. ohmeri, C. bracarensis, C. nivariensis, K. marxianus, P. norvegensis)*, followed by anidulafungin (2 species, *K. marxianus* and *P. norvegensis*), and caspofungin (1 species, *P*.*norvegensis*), which had more than two 2-fold dilution steps increasing of MIC at pH4.5 when compared with those at pH7.0.

Based on established ECV (≥0.5 mg/L) of *Cl. Lusitaniae* to micafungin, all nine isolates of *Cl. lusitaniae* have MIC_50_=0.125 mg/L, MIC_90_=0.25mg/L at pH7.0, while MIC_50_=0.5 mg/L, MIC_90_=1 mg/L at pH4.5 which are above the ECV. However, all nine isolates of *Cl. lusitaniae* had unchanged MICs to anidulafungin and caspofungin at both pH levels. Four isolates of *C. fabianii* had similar MIC as *Cl. lusitaniae*.

For two species of *C*.*bracarensis* (n=2), *C*.*nivariensi* (n=1) in *N. glabrata* complex, all 3 isolates were sensitive to micafungin at pH7.0, but 2 of them (MIC=0.5 mg/L) had MIC above the CBP of *N. glabrata* (0.25 mg/L) at pH4.5. However, the 3 isolates had unchanged MIC to anidulafungin and caspofungin at both pH levels, which Caspofungin MIC are 0.5mg/L (CBP of *N. glabrata* ≥0.5 mg/L), while anidulafungin MIC are < 0.5mg/L (CBP of *N. glabrata* ≥0.5 mg/L).

The one isolates of *P. norvegensis* shown susceptible to 3 echinocandins (MIC< 0.25 mg/L) at pH7.0, although it had two 2-fold dilution steps increasing of MIC to 3 echinocandins at pH4.5, only showing resistant to micafungin (CBP of *P. kudriavzevii*≥1 mg/L) at pH4.5.

Two isolates of *C. metapsilosis* had unchanged MIC to 3 echinocandins at two pH levels, one of the 2 isolates with MIC value above the ECV (0.25-1 mg/L).

*Diutina rugose* (n=1) and *Tricholoma matsutake* (n=1) had unchanged MIC with high MIC (0.25-2 mg/L) to 3 echinocandins at two pH levels.

For AmB, only two species (*K. ohmeri* and *D. rugosa*) had unchanged MICs at both pH levels, whereas the rest species had two 2-fold dilution steps increasing of MIC at pH4.5 which were >2 mg/L of MIC against the most yeast species, particular for *P. norvegensis* (n=1) with the most high MIC (8 mg/L) at pH4.5 when compared with those (MIC < 2 mg/L) at pH7.0. *D. rugosa*(n=1) showed high MIC (4/ 2 mg/L) while *K. ohmeri* (n=2) showed low MIC (0.5 mg/L) at both pH levels.

For nystatin, 7 species had two 2-fold dilution steps increasing of MIC at pH4.5 compared to those at pH7.0, and *P. norvegensis(*n=1)was the most affected by lower pH (8 mg/L) when compared to those (0.5 mg/L) at pH7.0.

All species displayed an unchanged MIC with a low MIC value (0.013-0.125 mg/L) to 5-flucytosine at two pH levels.

## Discussion

In our study, the main findings were the large variety of rare yeast species recovered in VVC. Based on recent phylogenetic relationships among *saccharomycotina* yeasts (11), 11 rare yeast species isolated from the vaginal samples belong to *metschnikowia* (*Cl. lusitaniae*), *wickerhamomycetaceae* (*C*.*fabianii*), *Kodamaea / Diutina* (*K. ohmeri, D. rugosa*), *Nakaseomyces* (*C. bracarensis, C. nivariensis*), *Lodderomyces* (*C. metapsilosis*), *Kluyveromyces* (*K. marxianus*), *Pichia* (*P. norvegensis*), *Tricholoma matsutake* and *C. africana* (n=1, respectively).

*Cl. lusitaniae* (n=9) was the most frequently identified species among 11 rare yeast species causing VVC in this study, followed by *Cy. fabianii* (n=4). Rest of each species contains 2 or1 isolates only. A study conducted in China on VVC displayed similar findings as ours regarding the isolation of rare yeast species (2). Indeed, this study conducted on 3574 yeast isolates obtained from patients with VVC identified 12 rare yeast species (< 0.1%, 20 isolates in total) based on molecular identification, which half of the species were presented in our study.

The antifungal agents most affected by acid pH were terconazole (4-fold higher) to 11 species; amphotericin B (3-fold higher) to 9 species except *K. ohmeri* and *D. rugose*; nystatin (2-fold higher) to 9 species except *C. nivariensis* and *C. africana*; micafungin (2-fold higher) to 8 species except *C. metapsilosis, D. rugosa* and *Tricholoma matsutake*, respectively (Table1-2, Table S3 3, Fig. 2-3). Our results are comparable to previous study by Spitze et al (10) that terconazole was the one affected the greatest by lower pH to *C. albicans, N. glabrata*, and 8 other yeast species. Previous studies (9, 10) had reported that activity of amphotericin B and nystain were profoundly affected by pH for *C. albicans* and *N. glabrata*, with a dramatic increase in the MIC_90_ at lower pH. Here for 11 rare yeast species tested, it displayed the same trend of high MICs for the rare yeast species to terconazole and amphotericin B, nystain as those for common yeast species.

Danby et al. (9) reported MIC_90_ values for 5-flucytosine were unchanged to *N. glabrata*, but a dramatic increased MIC were seen to *C. albicans* when tested at lower pH. In contrast, as observed in our study, 5-flucytosine was not affected by pH to all 11 tested species, and maintained activity at both pH levels.

Among 3 echinocandins tested in this study, caspofungin and anidulafungin were not affected significantly by lower pH, and showing activity against the most rare yeast species at both pH levels. Our results is similar as those reported by Danby (9) that caspofungin remained a stable MICs and activity against *C. albicans* and *N. glabrata* isolates with a decrease in pH. In contrast, an important new findings in the present study revealed that micafungin among 3 echinocandins was affected greatest by lower pH to 8 species except *C. metapsilosis, K. ohmeri* and *Tricholoma matsutake*, exhibited resistance to micafungin (MIC > 0.5-1mg/L) at pH4.5(Table 2, Table S3, Fig.3). Additional studies would need to be performed to evaluate echinocandins response *in vivo* as a topical compound.

*Cl. lusitaniae*, originally assigned to the teleomorphic state of *Candida lusitaniae*, belongs to the genus *Candida*/*Clavispora* within the family *Metschnikowiaceae* and the order *Saccharomycetales* (14). Nine isolates of *Cl. lusitaniae* exhibited low MIC values to all antifungals tested at pH7.0, while the acidic pH increased the MICs of several antifungals including posaconazole and terconazole among azoles, micafungin among 3 echinocandins, amphothericin B and nystatin(Table 1-2, Table S3, Fig.3), which are above the ECV for *Cl. Lusitaniae*(Table S2).

Previous studies reported that, *Cy. fabianii* has the potential to develop resistance to antifungals, including fluconazole, voriconazole, caspofungin, and amphotericin B (15). A similar observation in our study displayed an elevated MICs to poscanozole (MIC_50_/MIC_90_: 0.5/1 mg/L), miconazole (MIC_50_/MIC_90_: 1/2 mg/L), clotrimazole (MIC_50_/MIC_90_ 0.5/1 mg/L) at pH7.0, with significantly increased MIC at pH4.5 for terconazole (MIC_50_/MIC_90_: 4/8mg/L), amphotericin B (MIC_50_/MIC_90_: 4/4 mg/L), micafungin(MIC_50_/MIC_90_: 0.5/1 mg/L), nystatin (MIC_50_/MIC_90_: 4/8 mg/L) for 4 isolates of *Cy. fabianii* (Table 1-2, Table S3, Fig.3), when compared to those at pH7. Most of the data on the antifungal susceptibility of *K. ohmeri* isolates are derived from individual case reports. Since *K. ohmeri* exhibits variable susceptibility to fluconazole, in agreement with the literature(15), two isolate in our study showed an elevated MIC to posaconazole (0.125/0.25mg/L), one in 2 isolates also had higher MIC to fluconazole (8mg/L) at pH7.0, and it revealed an increased MIC to fluconazole (16 mg/L), micafungin (0.5 mg/L), terconazole (8 mg/L) and nystatin (8 mg/L) at pH4.5.Thus, rapid and accurate identification of *K. ohmeri*, as well as timely antifungal susceptibility testing are critical to enhance patient care.

*C. metapsilosis* is one of clinically relevant members of the *C. parapsilosis* complex, belong to azole and amphotericin B sensitive clade of *lodderomyces* as reported by Stavrou AA (11), two isolates of *C. metapsilosis* from vaginal tract were sensitive to all azoles tested, but resistant to amphotericin B (4 mg/L), less susceptibility to 3 echinocandins at both pH levels.

In our study, *C. bracarensis, C. nivariensis* which are phylogenetically closely related to *N. glabrata* were the species most affected by a lower pH to all azoles (Table 1-2, Fig.2). Both species show higher MIC values to all azoles at lower pH, but these isolates remained susceptible to all azole drugs tested compared by BCP or ECV of *N. glabrata*. However, it demonstrated a moderately higher MIC to terconazole (MIC > 4 mg/L). The significantly reduced susceptibility of *N. glabrata* to azoles at lower pH had been reported by Danby et al. (9) and spitze et al. (10).

*P. norvegensis*, a close relative of *P. kudriavzevii* (= *C. krusei*) in the *Pichia* clade(15) is known to exhibit high MICs to fluconazole(16), itraconazole and posaconazole (5, 17) and seem to be more frequently isolated from human samples (11, 18). In our study, one isolate of *P. norvegensis* displayed a significant influence by acidic pH with higher MIC value to fluconazole (32mg /L), 3 echinocandins (0.125-1mg /L) and nystatin (8mg /L), amphotericin B(8mg /L) at pH 4.5 than pH7.0 (Table 1-2, Table S3).

For the other species, it is hard to draw clear conclusions only with one isolates. Overall, our results demonstrated that the difference varies between antifungal agents and yeast species at pH4.5 than at pH7.0.

The limitations in our study is that the most rare species were tested only 1-2 isolates. Due to lack of the breakpoints for those rare yeast species, interpretation of these data and the clinical relevance of increased MICs are questionable. Further studies are necessary with more samples from different hospitals to investigate antifungal potency of rare yeast species causing VVC at acid pH.

## Conclusion

As demonstrated in the present study, the *in vitro* antifungal susceptibility at pH 7.0 did not consistently predict the activity of antifungals against those rare yeast species at vaginal pH environment. The noted shift toward rare yeasts in the clinic causes a challenge in therapeutic management due to their reduced antifungal susceptibility (11). Accurate identification of rare yeast isolates is crucial because some species are intrinsically resistant to widely used antifungals (19, 20).

## Data availability

GenBank accession numbers of D1/D2 sequences for 26 isolates studied are listed in Table S1.

## Ethics approval and Informed Consent

This study was approved by the Hospital Review Board and Ethics Committee of People’s Hospital, Suzhou New District, Suzhou, Jiangsu, China.

## Author Contributions

All authors listed, have made substantial, direct and intellectual contribution to the work, and approved the final manuscript for publication.

## Reference

1. Hillier SL, Austin M, Macio I, Meyn LA, Badway D, Beigi R. 2021. Diagnosis and Treatment of Vaginal Discharge Syndromes in Community Practice Settings. Clin Infect Dis 72:1538–1543.

2. Shi Y, Zhu Y, Fan S, Liu X, Liang Y, Shan Y. 2020. Molecular identification and antifungal susceptibility profile of yeast from vulvovaginal candidiasis. BMC Infect Dis 20:287.

3. Zhou Y, Deng, S., Zhang, L., Zhang, H., Zhao, R., Wang, Y., Sun, Y.,Zhu, H. 2023. Distribution of pathogenic yeast species of vulvovaginitis and its epidemiological characteristics. Chin J Mycol 18.

4. Bretagne S, Renaudat C, Desnos-Ollivier M, Sitbon K, Lortholary O, Dromer F, French Mycosis Study G. 2017. Predisposing factors and outcome of uncommon yeast species-related fungaemia based on an exhaustive surveillance programme (2002-14). J Antimicrob Chemother 72:1784–1793.

5. Guitard J, Angoulvant A, Letscher-Bru V, L’Ollivier C, Cornet M, Dalle F, Grenouillet F, Lacroix C, Vekhoff A, Maury E, Caillot D, Charles PE, Pili-Floury S, Herbrecht R, Raffoux E, Brethon B, Hennequin C. 2013. Invasive infections due to Candida norvegensis and Candida inconspicua: report of 12 cases and review of the literature. Med Mycol 51:795–9.

6. Agin H, Ayhan Y, Devrim I, Gulfidan G, Tulumoglu S, Kayserili E. 2011. Fluconazole-, amphotericin-B-, caspofungin-, and anidulafungin-resistant Candida ciferrii: an unknown cause of systemic mycosis in a child. Mycopathologia 172:237–9.

7. Sugita T, Takeo K, Ohkusu M, Virtudazo E, Takashima M, Asako E, Ohshima F, Harada S, Yanaka C, Nishikawa A, Majoros L, Sipiczki M. 2004. Fluconazole-resistant pathogens Candida inconspicua and C. norvegensis: DNA sequence diversity of the rRNA intergenic spacer region, antifungal drug susceptibility, and extracellular enzyme production. Microbiol Immunol 48:761–6.

8. Linhares IM, Summers PR, Larsen B, Giraldo PC, Witkin SS. 2011. Contemporary perspectives on vaginal pH and lactobacilli. Am J Obstet Gynecol 204:120 e1–5.

9. Danby CS, Boikov D, Rautemaa-Richardson R, Sobel JD. 2012. Effect of pH on in vitro susceptibility of Candida glabrata and Candida albicans to 11 antifungal agents and implications for clinical use. Antimicrob Agents Chemother 56:1403–6.

10. Spitzer M, Wiederhold NP. 2018. Reduced Antifungal Susceptibility of Vulvovaginal Candida Species at Normal Vaginal pH Levels: Clinical Implications. J Low Genit Tract Dis 22:152–158.

11. Stavrou AA, Lackner M, Lass-Florl C, Boekhout T. 2019. The changing spectrum of Saccharomycotina yeasts causing candidemia: phylogeny mirrors antifungal susceptibility patterns for azole drugs and amphothericin B. FEMS Yeast Res 19.

12. Ngai PV, Dat TH, Nhi LY, Linh TTK, Thu NT, Anh VL, Dung BTT, Tran PV, Hien NT, Son NT, Que TT, Anh DN. 2025. Distribution and antifungal susceptibility of Candida species causing vulvovaginal candidiasis and urinary tract infection in Medlatec healthcare system, Ha Noi city, Vietnam in 2023. Ther Adv Infect Dis 12:20499361241311465.

13. Oz Y, Gokbolat E. 2018. Evaluation of direct antifungal susceptibility testing methods of Candida spp. from positive blood culture bottles. J Clin Lab Anal 32.

14. Rodrigues de Miranda L. 1979. Clavispora, a new yeast genus of the Saccharomycetales. Antonie Van Leeuwenhoek 45:479–83.

15. Arastehfar A, Lass-Flörl C, Garcia-Rubio R, Daneshnia F, Ilkit M, Boekhout T, Gabaldon T, Perlin DS. 2020. The Quiet and Underappreciated Rise of Drug-Resistant Invasive Fungal Pathogens. J Fungi (Basel) 6.

16. Papon N, Courdavault V, Clastre M, Bennett RJ. 2013. Emerging and emerged pathogenic Candida species: beyond the Candida albicans paradigm. PLoS Pathog 9:e1003550.

17. Pfaller MA, Diekema DJ, Messer SA, Boyken L, Hollis RJ, Jones RN, International Fungal Surveillance Participant G. 2003. In vitro activities of voriconazole, posaconazole, and four licensed systemic antifungal agents against Candida species infrequently isolated from blood. J Clin Microbiol 41:78–83.

18. Pfaller MA, Diekema DJ, Gibbs DL, Newell VA, Ellis D, Tullio V, Rodloff A, Fu W, Ling TA, Global Antifungal Surveillance G. 2010. Results from the ARTEMIS DISK Global Antifungal Surveillance Study, 1997 to 2007: a 10.5-year analysis of susceptibilities of Candida Species to fluconazole and voriconazole as determined by CLSI standardized disk diffusion. J Clin Microbiol 48:1366–77.

19. Desnos-Ollivier M, Robert V, Raoux-Barbot D, Groenewald M, Dromer F. 2012. Antifungal susceptibility profiles of 1698 yeast reference strains revealing potential emerging human pathogens. PLoS One 7:e32278.

20. Whaley SG, Berkow EL, Rybak JM, Nishimoto AT, Barker KS, Rogers PD. 2016. Azole Antifungal Resistance in Candida albicans and Emerging Non-albicans Candida Species. Front Microbiol 7:2173.

